# *In silico* tools for accurate HLA and KIR inference from clinical sequencing data empower immunogenetics on individual-patient and population scales

**DOI:** 10.1101/2020.04.24.060459

**Authors:** Jieming Chen, Shravan Madireddi, Deepti Nagarkar, Maciej Migdal, Jason Vander Heiden, Diana Chang, Kiran Mukhyala, Suresh Selvaraj, Edward E. Kadel, Matthew J. Brauer, Sanjeev Mariathasan, Julie Hunkapiller, Suchit Jhunjhunwala, Matthew L. Albert, Christian Hammer

## Abstract

Immunogenetic variation in humans is important in research, clinical diagnosis and increasingly a target for therapeutic intervention. Two highly polymorphic loci play critical roles, namely the human leukocyte antigen (HLA) system, which is the human version of the major histocompatibility complex (MHC), and the Killer-cell immunoglobulin-like receptors (KIR) that are relevant for responses of Natural Killer (NK) cells and some subsets of T cells. Their accurate classification has typically required the use of dedicated biological specimens and a combination of *in vitro* and *in silico* efforts. Increased availability of next generation sequencing data has led to the development of ancillary computational solutions. Here, we report an evaluation of recently published algorithms to computationally infer complex immunogenetic variation in the form of HLA alleles and KIR haplotypes from whole-genome or whole-exome sequencing data. For both HLA allele and KIR gene typing, we identified tools that yielded >97% overall accuracy for 4-digit HLA types, and >99% overall accuracy for KIR gene presence, suggesting the readiness of in silico solutions for use in clinical and high-throughput research settings.

---

**T**he classical human leukocyte antigen (HLA) gene complex on chromosome 6, and the killer-cell immunoglobulin-like receptor (KIR) genes on the leukocyte receptor complex (LRC) on chromosome 19 are complex genomic loci that have been known to be difficult to genotype accurately. With the rapidly emerging treatment approaches in the fields of cancer immunotherapy^1,2^ and autoimmunity^3^, the accurate characterization of a patient’s immunogenetic composition in the HLA and KIR regions is becoming more clinically important.

HLA proteins play an important role in presenting peptides, lipids and glycolipids derived from self, tumor or microbial antigens. They display an extreme amount of allelic polymorphism, as a result of pathogen-driven and balancing selection.^4^ Much research has shown HLA variants to be strongly associated with multiple immune and non-immune phenotypes in the fields of cancer^5,6^, autoimmunity^3,7^, neurodegeneration^8^, and infectious diseases^3,7^. In the clinic, achieving matched HLA alleles between the donor and recipient is critical for organ and stem cell transplantation, such that HLA typing and matching have been integrated as part of standard clinical protocols for decades.^9–11^ HLA typing has also become increasingly important in diagnostics and clinical practice. For example, several approved drugs carry labels indicating increased risk for adverse events for carriers of specific HLA alleles.^12–14^

KIR proteins are receptors for HLA class I ligands, and are predominantly expressed on natural killer (NK) cells. In contrast to most HLA genes, the genes coding for KIR display extensive copy number polymorphism^15^, in addition to considerable allelic variation^16^ for each gene. KIR have shown significant associations with disease phenotypes, mainly in the fields of infectious diseases, autoimmunity, inflammatory diseases, and cancer.^17^ Associations were found for both single KIR genes, and when considering their interactions with specific HLA molecules. HLA-KIR interactions were demonstrated to predict the risk of organ rejection after kidney transplantation, suggesting a clinical use case for KIR typing.^18,19^ KIR proteins are also known to be important co-determinants of NK cell education, which is in part mediated through their interactions with different HLA molecules. Such interactions significantly define the heterogeneity of NK cell responsiveness and their sensitivity to inhibition by HLA across individuals.^20,21^ As such, KIR proteins play a critical role in the recognition of “missing-self” phenotypes in infected or tumor cells, which are typically defined by the loss or down-regulation of HLA class I cell surface expression.^22^

For research purposes, genotyping arrays covering single nucleotide polymorphisms (SNPs) across the genome have been used to impute HLA and KIR types. However, they require statistical imputation methods to disentangle the complex linkage disequilibrium (LD) between SNPs and HLA or KIR types.^23,24^ These methods also rely on the availability of ancestry-specific or multi-ancestry reference panels that can be difficult to obtain, especially for populations not well represented in genomic data sets.^25^ In clinical diagnostics, dedicated immunogenetics laboratory solutions to HLA and KIR genotyping are being continually developed.^26^ Initial molecular typing technologies were low throughput and/or probe-based assays.^27^ In recent years, high throughput next generation sequencing (NGS) has become increasingly affordable.^28^ This has enabled the prevalent use of whole-genome sequencing (WGS) and whole-exome sequencing (WES) in the clinic.^29,30^ While well-validated bioinformatics pipelines have been implemented to detect millions of genetic variants from the available clinical sequences,^31,32^ they are typically employed uniformly to the entire genome or exome, and can be ineffective at particular genomic loci that are highly polymorphic, such as the HLA and KIR regions. Dedicated *in silico* typing tools that use NGS data and specifically target the HLA or KIR genes could be a cost-effective and efficient alternative to traditional laboratory HLA or KIR typing methods. While such NGS-based approaches do not require linkage disequilibrium-based statistical imputation for genotyping (since the sequencing reads directly contain the information to define e.g. the HLA allele status), they do require the use of comprehensive databases that capture the diversity and complexity of these genomic loci for alignment (as opposed to single reference genomes).

Despite their biological significance and many practical advantages, the development of clinically-ready *in silico* HLA and KIR typing have thus far been largely hampered by the genetically complex and highly polymorphic nature of the two regions.^7,33^ Here, we conducted a survey of the current HLA and KIR typing capabilities for potential scaling and readiness in clinical applications. We evaluated available computational HLA inference tools by comparing the inferred HLA alleles from WES and WGS data to a gold standard dataset, which was generated using a commercial dedicated typing method. We also assessed and validated a recently published KIR method by Roe and Kuang^34^, which can be used to infer KIR gene presence and absence from WGS data.

## Methods

### Generation of a gold standard HLA reference dataset

The gold standard reference dataset contains 56 patient samples. Genomic DNA was extracted from 1 ml of EDTA whole blood on the Roche MagNA Pure 96 system using the MagNA Pure 96 DNA and Viral NA Large Volume Kit. They were then sent to LabCorp (Burlington, NC, USA) for sequence-based, two-field HLA genotyping, using accepted scientific standards meeting the accreditation requirements of the American Society for Histocompatibility and Immunogenetics (ASHI) and the College of American Pathologists (CAP). HLA class I genotypes of *HLA-A, -B*, and *-C* were determined using a combination of long-range sequencing from PacBio (Menlo Park, CA, USA) RSII and Sanger sequencing. Class II genotypes for *HLA-DPA1, -DPB1, -DQA1, -DQB1, -DRB1, -DRB3, -DRB4, -DRB5* were determined using a combination of long-range sequencing from PacBio RSII, and NGS from Illumina (San Diego, CA, USA) TruSight and MiSeq technologies.

### Generation of a KIR reference dataset

Genotyping of KIR alleles was performed for 72 patients using the LinkSeq KIR kit according to manufacturer’s user guide (Catalog No: 5358R, One Lambda, Canoga Park, CA, USA). Briefly, the human genomic DNA was amplified and melt curves were collected on a real-time PCR instrument (QuantStudio 5 system, Thermo Fisher Scientific, Waltham, MA, USA). The data was exported to SureTyper software 6.1.2 (One Lambda) for interpretation and reporting of the genotype.

### Whole-genome and whole-exome sequencing

WGS and WES were performed for the samples in the HLA and KIR reference datasets, on Illumina HiSeq instruments using paired-end reads of 150 bp. Each whole genome (WG) was sequenced at an average of 30x coverage. For WES, all samples passed QC criteria of 85% of targeted bases at 20x or greater coverage (+/- 5%). FASTQ files were generated for each WG and exome, and were used as direct inputs for each tool, where appropriate. For tools that required BAM files as inputs, the FASTQ files were processed according to a workflow built using the GATK best practices from 2015 and GATK v3.5^32^. Briefly, it includes read alignment using bwa 0.7.15 with GRCh38 as the reference genome, duplicate marking using Picard tools v2.9, indel re-alignment using GATK v3.5, and then base quality score recalibration using GATK 3. For bwa, except for the ‘-M’ flag, all the default options were used. Individual-level genetic data for this study cannot be made publicly available due to consent restrictions.

### Selection and configuration of HLA typing tools

For WGS and WES HLA genotyping, we specifically selected tools that (1) are recently published, (2) are easy to install and implement, (3) have the ability to work with WGS and WES data, (4) and could genotype both HLA I and II genes for potential clinical use. These included HLA*PRG:LA,^35^ xHLA v1.2 [https://github.com/humanlongevity/HLA]^36^, HLA-HD v1.2 [https://www.genome.med.kyoto-u.ac.jp/HLA-HD]^37^, and HISAT2 v2.1 [https://ccb.jhu.edu/software/hisat2/index.shtml]^38^. As a comparison to a more widely used tool, we also assessed results from Polysolver v1.0^39^. Polysolver has been widely utilized in many benchmarking and research studies, and has been shown to be one of the more accurate HLA class I genotyping tools.^40-44^ Polysolver requires hg18 or hg19 as the human reference genome for read extraction from chromosome 6 in the BAM files; to overcome this limitation, we modified the code to include the use of BAMs that are aligned to hg38 as well. No other modifications were made to the code. For xHLA, BAMs were aligned to hg38, and then preprocessed with an additional bash script provided by the authors [https://github.com/humanlongevity/HLA/blob/master/bin/get-reads-alt-unmap.sh]. For HLA-HD and HISAT2, the raw FASTQs were used directly. All the tools were run using default settings, and with the HLA/IMGT databases versions that came with the respective tools. For the exact run parameters, please refer to Supplementary Table 1.

### Evaluation of HLA typing tools

The tools typically give two alleles per HLA gene, but we do see in occasions, albeit rare, in our study where the algorithm provided more than one pair of possible alleles, and often in descending order of significance. For any results that offer more than two alleles, we only took the top two inferred HLA alleles; we assumed a diploid germline genome.

Each *in silico* HLA typing tool was evaluated by the number of concordant HLA alleles called by the tool when compared with the reference HLA genotypes obtained from LabCorp, for the classical HLA I and II genes. Its accuracy was defined as the quotient of the number of concordant calls and the sum of the number of concordant plus discordant calls. For uniformity, results from the tools were converted to four-digit resolution before evaluation, except for HLA*PRG:LA, which can only perform genotyping at G group resolution. A G group contains HLA alleles that have the same genomic sequence for the same binding site, while a P group contains all the HLA alleles that have the same protein sequence for the antigen binding site of the HLA binding molecule (exons 2 and 3 for HLA class I and exon 2 for class II). A minimum of P group resolution or higher (including G group, four-digit/ two-field resolutions), is usually considered ‘high resolution typing’ and therefore clinically relevant.^45^ For HLA*PRG:LA, gold standard results were first converted to G groups before evaluation. G group information was obtained from the IMGT/HLA database [http://hla.alleles.org/alleles/g_groups.html].

We split the evaluation of the tools’ accuracy into three categories: (1) HLA I, consisting of classical HLA I genes *HLA-A, -B*, and *-C*, (2) HLA IIa, consisting of classical HLA II genes *HLA-DPA1, -DPB1, -DQA1, -DQB1*, and *-DRB1*, and (3) HLA IIb, consisting of a second group of classical HLA II genes, *HLA-DRB3, -DRB4*, and *-DRB5* (Table 1). We created a second category (HLA IIb) for additional DR genes because only HLA-HD currently genotypes them. Additionally, since every individual carries a variable copy number of the three genes *HLA-DRB3, -DRB4* or -*DRB5* that is highly dependent on the -*DRB1* genotype,^46^ a no-call by the computational tool is considered concordant with the gold standard results if it is not also identified. Some tools cannot genotype the full set of the classical class II genes, thus we also provided the accuracy with respect to each class II gene (Table 2).

**Table 1.**
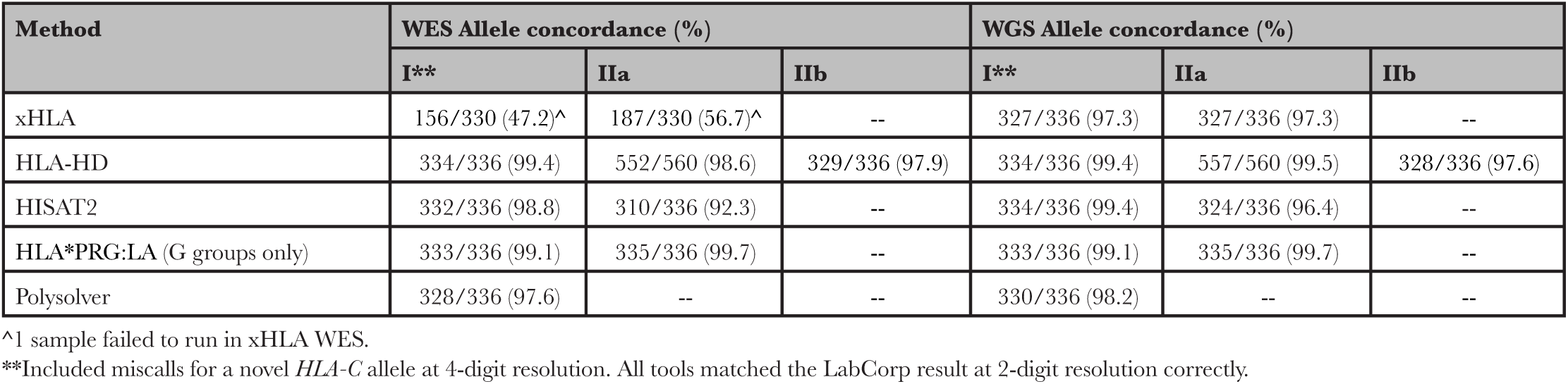
Overall evaluation results for HLA typing using WES and WGS. HLA genes are categorized into classical class I genes (*A, B, C*), IIa genes (*DPA1, DPB1, DQA1, DQB1, DRB1*), and IIb genes (*DRB3, DRB4, DRB5*). The class IIa genes that each tool can genotype differ: HLA-HD can infer all of the above; xHLA can only infer *DPB1, DQB1* and *DRB1*; HISAT2 and HLA*PRG:LA only *DQA1, DQB1* and *DRB1*. Note that HLA*PRG:LA was evaluated based on the G group resolution. Results from the older and well-utilized Polysolver were provided as an additional source of comparison for HLA I genes.

**Table 2.** Evaluation results by HLA II genes for HLA typing using WES and WGS. Note that HLA*PRG:LA was evaluated based on the G group resolution, not at 4-digit resolution.

| HLA II gene | xHLA (%) |  | HLA-HD (%) |  | HISAT2 (%) |  | HLA*PRG:LA (%) |  |
| --- | --- | --- | --- | --- | --- | --- | --- | --- |
|  | WES<sup>^</sup> | WGS | WES | WGS | WES | WGS | WES | WGS |
| *DPA1* | -- | -- | 112/112 (100) | 112/112 (100) | -- | -- | -- | -- |
| *DPB1* | 70/110 (63.6) | 109/112 (97.3) | 110/112 (98.2) | 110/112 (98.2) | -- | -- | -- | -- |
| *DQA1* | -- | -- | 106/112 (94.6) | 111/112 (99.1) | 111/112 (99.1) | 110/112 (98.2) | 112/112 (100) | 112/112 (100) |
| *DQB1* | 53/110 (48.2) | 107/112 (95.5) | 112/112 (100) | 112/112 (100) | 111/112 (99.1) | 110/112 (98.2) | 111/112 (99.1) | 112/112 (100) |
| *DRB1* | 64/110 (58.2) | 111/112 (99.1) | 112/112 (100) | 112/112 (100) | 88/112 (78.6) | 104/112 (92.9) | 112/112 (100) | 111/112 (99.1) |
| *DRB3* | -- | -- | 110/112 (98.2) | 111/112 (99.1) | -- | -- | -- | -- |
| *DRB4* | -- | -- | 110/112 (98.2) | 108/112 (96.4) | -- | -- | -- | -- |
| *DRB5* | -- | -- | 109/112 (97.3) | 109/112 (97.3) | -- | -- | -- | -- |

<sup>^</sup>1 sample failed to run in xHLA WES.

### Evaluation of KIR typing with kpi and interpretation of results

For inference of KIR gene presence or absence, we evaluated kpi [https://github.com/droeatumn/kpi, downloaded March 11^th^, 2020]. An earlier version of the software was recently presented in a preprint and did not provide a validation of its accuracy when compared to qPCR-based dedicated KIR typing methods.^34^ Kpi requires WGS FASTQ files as input data and outputs a presence / absence call for each KIR gene, as well as possible combinations of KIR haplotypes according to a provided list of reference haplotypes [https://github.com/droeatumn/kpi/blob/master/input/haps.txt]. Each KIR gene can in principle be characterized by copy number and allelic variation. A KIR haplotype determines the order and presence of single KIR genes.^47,48^ However, kpi only detects presence or absence of each KIR gene, not allele status or copy number. As a result, the calls for haplotype pairs can be ambiguous (due to differences in copy number of present genes), but the presence of single KIR genes can be resolved.^49^ As such, KIR typing with kpi, albeit coarse, is still useful because many associations have been reported on haplotype or gene level. Furthermore, interactions of KIRs with their HLA ligands are usually defined at the KIR gene level.^17,22^ It should be noted though, that the KIR genes do show extensive allelic polymorphism that can still have an effect on such defined interactions.^33,50^

### Inference of HLA-KIR interactions

NK cell inhibiting as well as activating KIR interactions with their HLA class I ligands were defined according to Pende *et al*.^18^ Briefly, some KIR interact with groups of 4-digit HLA alleles according to specific HLA amino acid residues. *HLA-B* alleles were classified as either *Bw4* or *Bw6* according to amino acid positions 77 - 83. *HLA-C* alleles were assigned C1 or C2 status based on amino acid position 80.^18^ Other interactions were defined between KIR and specific 2-digit or 4-digit HLA alleles (e.g. *A*03 - KIR3DL2*).

### Data availability

Owing to consent restrictions, individual-level genetic data used for this study cannot be made available.

## Results

### High accuracy for HLA I and II typing with current gold standard WES and WGS data

We selected HLA genotyping tools to infer HLA identities of 56 patients from the EXCELS (NCT00252135)^51^ and AVANT (NCT00112918)^52^ clinical trials. The inferred HLA types were then compared to results from the gold standard reference dataset. Despite the limited size of our dataset, the diversity of HLA class I alleles for each HLA gene in our samples is highly comparable to the publicly available, ethnically varied and sequencing-derived 1000 Genomes/HapMap validation set generated by Ehrlich *et al*.^53^ (Supplementary Table 2).

From the results consolidated in Table 1, all the selected genotyping tools perform generally well, at an accuracy of >90% for most of the class I and II gene categories, except for xHLA on WES class I and IIa genes. xHLA demonstrated uncharacteristically low accuracy for our WES data for both classes I and II, when compared to both reported performance^36^ and the other tools. HISAT2 and HLA-HD performed comparably for the HLA I genes *A, B* and *C* for both WES and WGS data, at >98.8% accuracy. Overall, for class II genes, HLA-HD is consistently the most accurate HLA typing tool for both WES and WGS data, at >97.6% accuracy for *DRB3, DRB4*, and *DRB5*, and >98.6% accuracy for the *DPA1/B1, DQA1/B1* and *DRB1* genes. Moreover, it also provides the widest range of HLA II genes, with the ability to genotype all the classical class II genes (including *HLA-DRA, -DRB3, -DRB4* and *-DRB5*), while other tools are restricted to a subset of classical class II alleles. However, HLA-HD has lower accuracy for *DQA1* when working off WES data, compared to HISAT2 (Table 2).

While we observed similar or increased accuracy when comparing results from WGS to WES data from the same tool (Table 1), WGS and WES miscalls were not always the same. This is evident when assessing accuracy at the gene level. For example, the increase in overall accuracy of HISAT2 when using WGS data for identifying class II alleles was mainly due to a lower *HLA-DRB1* accuracy when using WES data.

We next focused on the top performant methods, HLA-HD and HISAT2. Notably, the number of miscalls in HLA I genes was too low to characterize patterns or biases in the miscalls (Table 1; maximum of 4 miscalls). For class II genes, HISAT2 showed the highest number of *HLA-DRB1* miscalls when using WES data (Table 2). By contrast, HLA-HD miscalled mostly in the class IIb genes, where it failed to discriminate the highly similar alleles of the paralogous genes, *HLA-DRB3, -DRB4* and *-DRB5*. There was no obvious bias in miscalls of homozygous genes in the HLA-HD and HISAT2 results (Supplementary Table 3). We note that most of HLA-HD miscalls still called an allele in the same G group as the anticipated allele. In particular, almost all the miscalls in *HLA-DQA1* (5/6 of -*DQA1* miscalls and 5/8 of total miscalls) were made when they were called as -*DQA1*03:01* and the anticipated calls were *-DQA1*03:02* and -*DQA1*03:03*; all three alleles are in the same G group. For HISAT2, many of the miscalls in class II genes were due to missing calls, i.e. calls that the tool was not able to assign an allele at all.

### High accuracy for identifying KIR gene absence/presence

We used kpi to infer KIR gene presence for 824 patients with available WGS data from the AVANT trial (NCT00112918).^52^ We found that the gene carrier frequencies were very similar to those of published KIR gene carrier frequencies from an English cohort of Caucasoid ethnicity.^54^ (Figure 1a). The software was unable to predict possible haplotype pairs for 3% (n=25) of cases.

**Figure 1.**
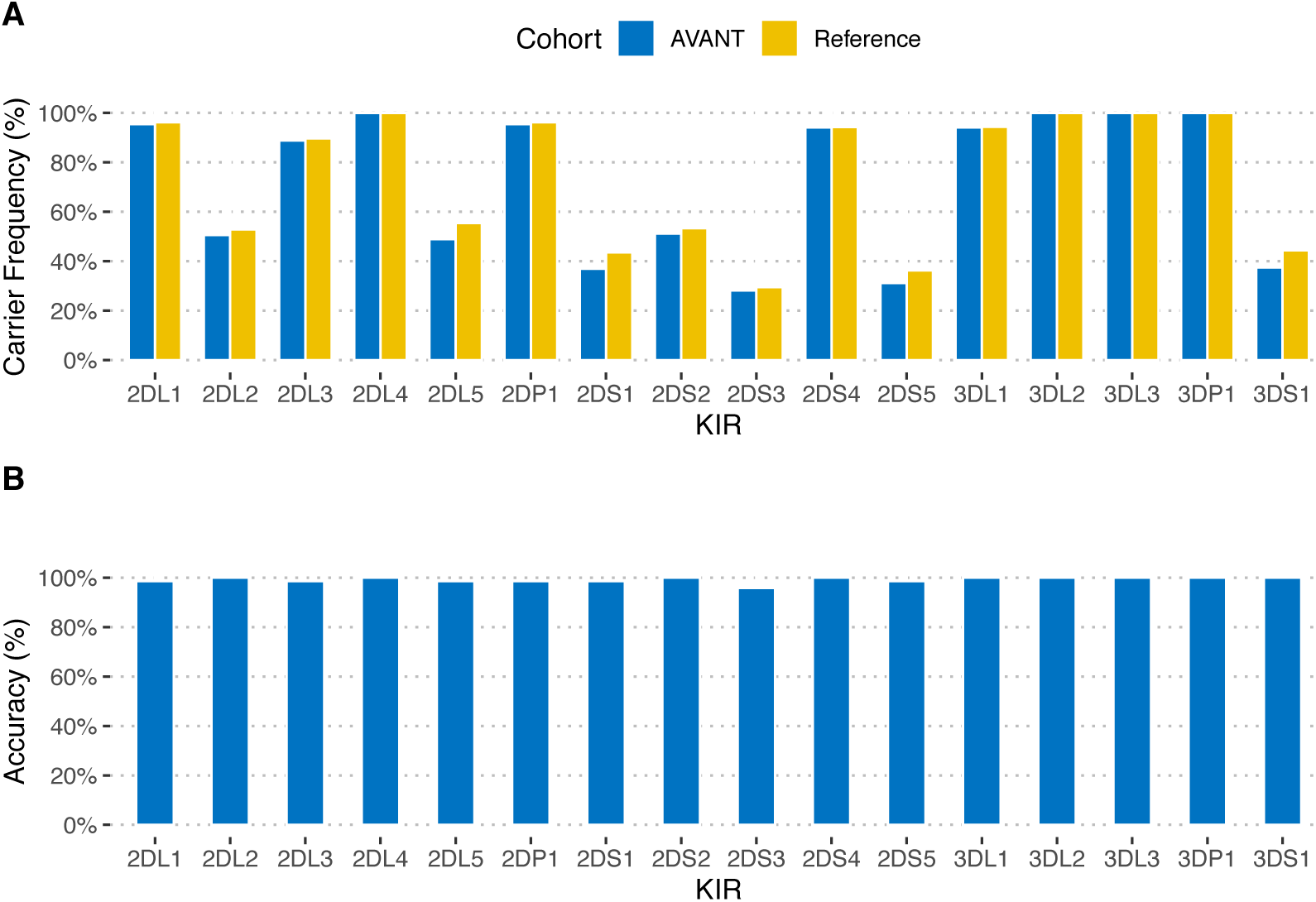
KIR gene carrier frequencies and accuracy of kpi typing. **(A)** KIR gene presence for AVANT patients (N=824) was inferred from kpi haplotype predictions, and compared to published frequencies for an English reference cohort (N=584). **(B)** For 72 AVANT patients typed with kpi, KIR typing with a qPCR-based method (LinkSeq) was performed to assess typing accuracy.

We selected 72 of these 824 patients to perform qPCR-based KIR typing, based on DNA sample availability and the diversity of kpi-predicted KIR haplotypes. These patients are different from those selected for the HLA gold standard data because of limited availability of DNA. For this selection, we also included eight of the 25 patients that had yielded uninterpretable haplotype combinations using kpi.

99.2% of kpi gene presence/absence calls were correct when compared to our qPCR-based reference (range of 95.8% -100% for the 16 tested genes, Figure 1b, Supplementary Table 4). When excluding the four framework genes *KIR3DL3, KIR3DP1, KIR2DL4*, and *KIR3DL2*, which were invariable in our dataset and are present in most common haplotypes, the accuracy for the 12 remaining genes was 99.0%. Three patients were typed with one error each (*KIR2DS3*), and one patient with six errors. The eight qPCR-typed individuals with uninterpretable kpi haplotype results had correct kpi gene calls, which could not be clearly assigned to a reference haplotype combination as provided by the software (Supplementary Table 5).

## Discussion

The broad availability of NGS data, generated in a multitude of clinical scenarios, allows for the inference of disease-relevant immunogenetic variation without additional dedicated typing efforts. Hence, in order to evaluate the usefulness in a clinical setting, we were interested in a systematic comparison of the newest generation of computational HLA typing solutions, run alongside the more well-established HLA typing tool Polysolver, that is limited in only typing HLA I genes. Our analyses suggest that for both WES and WGS data, most of the tools outperform Polysolver in HLA inference, in both class I genotyping accuracy and the ability to perform inference on HLA II genes. For KIR typing using kpi, we are not aware of a published independent validation of its performance. Our evaluation demonstrates that kpi performs well at determining the absence or presence of a KIR gene, but it is not able to ascertain KIR allele or gene copy number.

Our analyses also show the breadth of class II genes that current state-of-the-art tools can infer. In particular, only HLA-HD was able to genotype the classical HLA II genes, *HLA-DRB3*, -*DRB4*, and -*DRB5*, and we were able to further examine the results with our gold standard dataset. Interestingly, we found that many of the *HLA-DRB3, -DRB4* and *-DRB5* miscalls can be salvaged using knowledge of the strong (and clear) linkage disequilibrium between the *HLA-DRB1* gene and its *DRB* paralogs.^55^ It appears that incorporating this piece of biological information could be useful in developing tools that would like to genotype all the *DRB* genes, especially when there is high accuracy in genotyping the *DRB1* gene.

Additionally, with WGS and WES data for the same subjects, we observed that HLA inference from WGS data has yielded marginally higher accuracy compared to WES in many HLA genes (Tables 1 and 2). This possibly indicates that the addition of non-coding sequence information, or a more uniform read coverage in the HLA region, might be more relevant in these genes, especially in resolving alleles that are in the same G group, e.g. *HLA-DQA1*03:01*, -*DQA1*03:02* and -*DQA1*03:03*. It might also point to the use of bait sets in WES, which can bias the calling of some alleles; WGS does not require such baits.

Of note, the dedicated HLA typing approach (LabCorp) identified one novel *HLA-C* allele in our cohort of 56 individuals at two-digit resolution. Even though all the tools gave an estimation (i.e. it was not a missing call) and were correct at the two-digit resolution, none of the tested tools were able to identify the allele as novel. This is because the inferences are all based on aligning sequencing reads to a database of known HLA alleles. This might not be highly important for large-scale genetic association approaches, but might be relevant in a clinical setting focused on individual patients, especially for ethnicities that have thus far been underrepresented with regard to genome sequencing, let alone HLA typing.^56,57^

The choice and accuracy of a given HLA method might depend on read length and sequencing coverage, which are factors that are not included in the current study. A recent comparison of HLA-HD and xHLA for use with target capture methods or amplification sequencing suggested that HLA-HD might decrease in sensitivity at read lengths below 150 base pairs (paired-end).^58^

We noticed that documentation for most of the HLA typing tools tested is mainly centered on the final inference, but not the auxiliary output files. The latter set of files typically contain the scores for all the candidates used for inference. While accuracy is important, tool documentation in a clinical setting is also imperative to better understand the tool and its outputs, so that best practices can be developed in the clinic for different contexts.

As for the KIR typing efforts, kpi was shown to predict KIR gene presence/absence at >99% accuracy overall, and at >95% for each gene. Six out of nine errors were found in a single individual. This was likely due to a sample swap, and the remaining three miscalls were all for *KIR2DS3* in different patients. In this case, all other genes were inferred at 100% accuracy. However, since kpi detects gene presence/ absence and does not perform an estimation of copy number, it assigns one or more possible haplotype combinations, resulting in considerable ambiguity. Thus, we recommend to analyze kpi results at the level of individual KIR genes, if possible. It is likely that the 25 uninterpretable haplotype pairs are due to carriers of rare haplotypes not present in the reference, which would prevent an assignment of possible reference haplotype combinations.

Notably, kpi requires WGS data and does not consider allelic variation within KIR genes. This is a significant limitation, since allotypes for a given KIR gene can be functionally different,^59^ and also have differential binding capacities to their predicted HLA ligands.^50,60,61^ Allele-level typing would be desirable. The only software presently available that we are aware of that provides this level of granularity was not designed to work with NGS data in a high-throughput setting.^62^

In conclusion, our survey for both high-resolution four-digit clinically-relevant HLA typing and inference of KIR gene presence from NGS data (of conventional read length and coverage) indicated that recently published software tools can yield very high accuracy (>97% for HLA alleles and >95% for KIR genes, respectively) that may be suitable not only for research us, but also for the clinic. For comparison, the 2019 Standards for Accredited Laboratories issued by the American Society for Histocompatibility and Immunogenetics only requires a minimum of 80% concordance with another CLIA-certified ASHI-accredited laboratory to be deemed satisfactory in clinical testing.^63^ It is noteworthy that WGS and WES continue to become less expensive, thereby presenting an alternative even in scenarios that focus only on HLA or KIR typing. However, a foreseeable hurdle is the process of obtaining regulatory approval for computational tools for HLA and KIR typing in the clinic, either as a stand-alone device, or as part of a pipeline. Such a process could be tricky as it can be highly dependent on the context of how the tool is being applied in the clinic. There are pros and cons for each tool. Other considerations in the choice of method that we did not explore in this study and might merit investigation in the future, include the characteristics of the NGS datasets at hand, such as the read length and read coverage, which can affect accuracy and thus cause deviations from what is shown in the present report.

Finally, we would like to further emphasize that computational tools can generate HLA and KIR information in a high-throughput manner on large cohorts of patients with clinical sequencing. Furthermore, the time and logistical challenges and risks associated with acquisition, preparation and shipping of valuable clinical specimens to perform a separate genotyping would be greatly reduced. In a clinical setting, the HLA and KIR results from these tools can then be applied directly to detect immunogenetic biomarkers that might be relevant for treatment decisions, or to predict the likelihood of adverse events for a given treatment of choice.^64^ HLA typing is also a requirement in the context of individualized cancer treatment strategies, including immunization efforts and neoadjuvant-directed T-cell therapies.^65,66^ Neoepitope prediction requires highly accurate HLA types in order to maximize the likelihood of an immunogenic anti-tumor response.^67^ KIR genes are emerging biomarkers in several disease areas, including cancer immunology,^68^ and should ideally be investigated in the context of their interactions with their HLA ligands. Having both HLA and KIR information will also allow stratification of patients according to their individual and biologically relevant HLA-KIR interactions (Supplementary Figure 1).^22^ As more computational tools for HLA and KIR typing are likely being developed in the future, they should be continuously evaluated so that they can fulfill a greater role in assessing clinical genomes.

## Acknowledgments

The authors would like to thank the author of kpi, David Roe, for helpful discussions. We are also thankful to Sean Kelley for support of this study.

## Notes

### Competing Interest Statement

JC, SM, DN, MM, JVH, DC, KM, SS, EEK, SM, JH, SJ, and CH are employees of Roche/Genentech. MJB is an employee of Maze Therapeutics. MLA is an employee of insitro.

