## Supplementary Data for "*In silico* tools for accurate HLA and KIR inference from clinical sequencing data empower immunogenetics on individual-patient and population scales"

### Table of Contents


| | |
|---|---|
| **Supplementary Table 1.** Selection and specification of HLA typing tools | 2 |
| **Supplementary Table 2**: Allele diversity of HLA and KIR genes in gold standard dataset | 3 |
| **Supplementary Table 3**: No obvious biases in HLA-HD and HISAT2 miscalls due to alleles that are homozygous or possessing the same peptide binding region (G groups) | 4 |
| **Supplementary Table 4**: Overall evaluation results for KIR typing from WGS data with kpi | 5 |
| **Supplementary Table 5**: qPCR-typed individuals with uninterpretable kpi results. | 6 |
| **Supplementary Figure 1**: Frequencies of HLA-KIR pairs with known biological relevance | 7 |

**<u>Supplementary Table 1</u>. Selection and specification of HLA typing tools**

| HLA typing tool | Configuration |
|---|---|
| xHLA | *Preprocessing (read extraction)*<br>get-reads-alt-unmap.sh S1_input.bam S1_input_pp.bam hg38.chr6.fna<br><br>*xHLA run*<br>run.py --sample_id S1 --input_bam_path S1_input_pp.bam --output S1_output |
| HLA-HD | hlahd.sh -t 4 -m 100 -f hlahd/freq_data S1_1.fastq S1_2.fastq hlahd/HLA_gene.split.txt hlahd/dictionary S1 S1_hlahd_output |
| HISAT2 | *Preprocessing (read extraction)*<br>hisatgenotype_extract_reads.py --base hisat2/index/genotype_genome --read-dir fastq/S1 --out-dir S1_extracted_reads<br><br>*HISAT2 run*<br>hisatgenotype.py --base hisat2/index/genotype_genome --region-list hla.A,hla.B,hla.C,hla.DQA1,hla.DQB1,hla.DRB1 --assembly -p 4 -1 S1_extracted_reads/S1.hla.extracted.1.fq.gz -2 S1_extracted_reads/S1.hla.extracted.2.fq.gz |
| HLA*PRG:LA | inferHLATypes.pl --BAM S1.bam --graph PRG_MHC_GRCh38_withIMGT --sampleID S1 --maxThreads 8 --outdir S1_HLA_LA_output |
| Polysolver | shell_call_hla_type S1.bam Unknown 1 hg38 STDFQ 0 S1_output |

Parameters used in the commands for the various HLA tools. The default arguments were used in each tool for both WES and WGS runs, including the IMGT version that came with the tool, unless specified otherwise. Note that for HLA*PRG:LA, a graph was built with default settings prior to this step.

**<u>Supplementary Table 2</u>: Allele diversity of HLA and KIR genes in gold standard dataset**

| | **1000 Genomes Project/Hapmap*** | **Gold standard in this study** |
|---|---|---|
| Number of unique samples | 183 | 56 |
| Number of HLA alleles | | |
| *A* | 34 | 23 |
| *B* | 54 | 40 |
| *C* | 27 | 28 |
| *DPA1* | -- | 4 |
| *DPB1* | -- | 20 |
| *DQA1* | -- | 12 |
| *DQB1* | -- | 16 |
| *DRB1* | -- | 26 |
| *DRB3/4/5* | -- | 4/2/3 |

* Samples that were HLA-genotyped from the 1000 Genomes Project in Erlich *et al.* (2011) contain replicates (253 samples, including replicates)

Diversity of HLA alleles in each HLA gene is comparable in the publicly available 1000 Genomes Project and the gold standard dataset used in this study.

**<u>Supplementary Table 3</u>: No obvious biases in HLA-HD and HISAT2 miscalls due to alleles that are homozygous or possessing the same peptide binding region (G groups)**

| | | HLA-HD | | HISAT2 | |
|---|---|---|---|---|---|
| | | **WES** | **WGS** | **WES** | **WGS** |
| Number of miscalls where the correct call is homozygous | HLA I^ | 0/2 | 0/2 | 0/4 | 0/2 |
| | HLA IIa | 2/8 | 0/3 | 2/26 | 4/12 |
| Number of miscalls where the correct call is in the same G or P group | HLA I^ | 1/2 | 1/2 | 1/4 | 1/2 |
| | HLA IIa | 8/8 | 2/3 | 0/26 | 0/12 |

^ Note that the miscalls included the novel *HLA-C* allele (called at 2-digit resolution by LabCorp)

**<u>Supplementary Table 4</u>: Overall evaluation results for KIR typing from WGS data with kpi**

| KIR gene | % accuracy |
|---|---|
| 3DL3 | 100.00% |
| 2DS2 | 100.00% |
| 2DL2 | 100.00% |
| 2DL3 | 98.61% |
| 2DP1 | 98.61% |
| 2DL1 | 98.61% |
| 3DP1 | 100.00% |
| 2DL4 | 100.00% |
| 3DL1 | 100.00% |
| 3DS1 | 100.00% |
| 2DL5 | 98.59% |
| 2DS3 | 95.83% |
| 2DS5 | 98.61% |
| 2DS4 | 100.00% |
| 2DS1 | 98.61% |
| 3DL2 | 100.00% |

For 72 AVANT patients typed with kpi, KIR typing with a qPCR-based method (LinkSeq) was performed to assess typing accuracy. The LinkSeq approach failed in 1 case for KIR2DL5, and this case was removed from the comparison for this gene.

**<u>Supplementary Table 5</u>: qPCR-typed individuals with uninterpretable kpi results.**

| ID | 3DL3 | 2DS2 | 2DL2 | 2DL3 | 2DP1 | 2DL1 | 3DP1 | 2DL4 | 3DL1 | 3DS1 | 2DL5 | 2DS3 | 2DS5 | 2DS4 | 2DS1 | 3DL2 |
|---|---|---|---|---|---|---|---|---|---|---|---|---|---|---|---|---|
| uninterp1 | 1 | 0 | 0 | 1 | 1 | 1 | 1 | 1 | 1 | 1 | 1 | 1 | 1 | 1 | 1 | 1 |
| uninterp2 | 1 | 0 | 0 | 1 | 1 | 1 | 1 | 1 | 1 | 0 | 0 | 0 | 0 | 1 | 1 | 1 |
| uninterp3 | 1 | 1 | 1 | 0 | 0 | 0 | 1 | 1 | 1 | 0 | 1 | 1 | 0 | 1 | 1 | 1 |
| uninterp4 | 1 | 1 | 1 | 0 | 1 | 1 | 1 | 1 | 1 | 0 | 1 | 1 | 1 | 1 | 1 | 1 |
| uninterp5 | 1 | 1 | 1 | 1 | 1 | 1 | 1 | 1 | 1 | 0 | 1 | 0 | 1 | 1 | 1 | 1 |
| uninterp6 | 1 | 1 | 1 | 0 | 0 | 0 | 1 | 1 | 1 | 0 | 1 | 0 | 1 | 1 | 1 | 1 |
| uninterp7 | 1 | 1 | 1 | 1 | 1 | 1 | 1 | 1 | 1 | 0 | 1 | 1 | 0 | 1 | 1 | 1 |
| uninterp8 | 1 | 0 | 0 | 1 | 1 | 1 | 1 | 1 | 1 | 1 | 1 | 1 | 0 | 1 | 0 | 1 |

qPCR-typed individuals with uninterpretable kpi results had gene presence / absence patterns that could not be clearly assigned to a reference haplotype combination.

**<u>Supplementary Figure 1</u>: Frequencies of HLA-KIR pairs with known biological relevance**

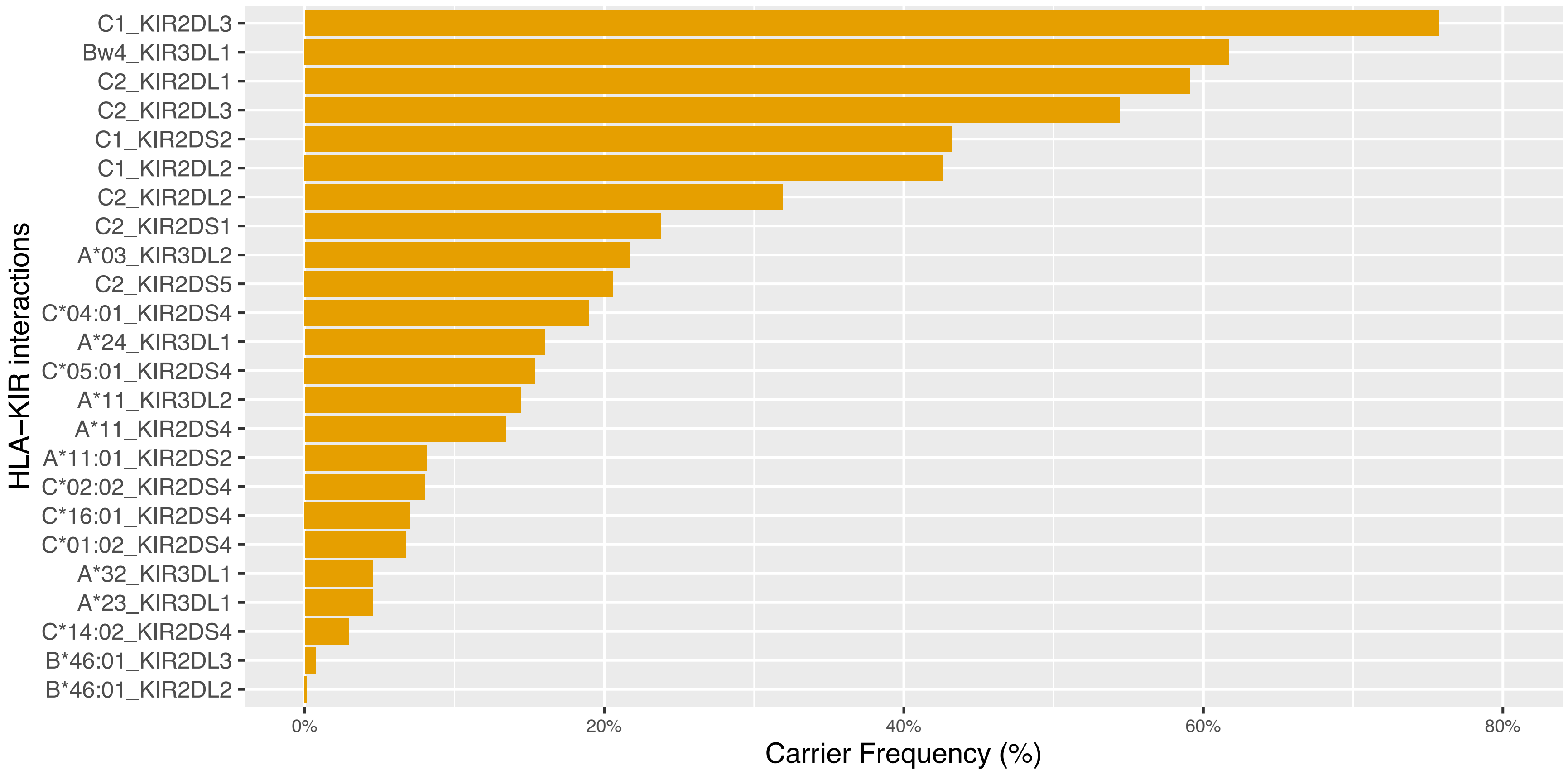


Experimentally validated HLA-KIR interactions for 824 AVANT patients were coded according to Pende *et al*[22].
